# Palmitic acid induces inflammation in placental trophoblasts and impairs their migration toward smooth muscle cells through plasminogen activator inhibitor-1

**DOI:** 10.1101/2020.05.13.085076

**Authors:** Amanda M. Rampersaud, Caroline E. Dunk, Stephen J. Lye, Stephen J. Renaud

**Author notes:** To whom correspondence can be addressed: Stephen J Renaud, Department of Anatomy and Cell Biology, University of Western Ontario, 1151 Richmond St, London, Ontario, Canada, N6A5C1.

## Abstract

A critical component of early human placental development includes migration of extravillous trophoblasts (EVTs) into the decidua. EVTs migrate toward, and displace vascular smooth muscle cells (SMCs) surrounding several uterine structures, including spiral arteries. Shallow trophoblast invasion features in several pregnancy complications including preeclampsia. Maternal obesity is a risk factor for placental dysfunction, suggesting that factors within an obese environment may impair early placental development. Herein, we tested the hypothesis that palmitic acid, a saturated fatty acid circulating at high levels in obese women, induces an inflammatory response in EVTs that hinders their capacity to migrate toward SMCs. We found that SMCs and SMC-conditioned media stimulated migration and invasion of an EVT-like cell line, HTR8/SVneo. Palmitic acid impaired EVT migration and invasion toward SMCs, and induced expression of several vasoactive and inflammatory mediators in EVTs, including endothelin, interleukin (IL)-6, IL8, and PAI1. PAI1 was increased in plasma of women with early-onset preeclampsia, and PAI1-deficient EVTs were protected from the anti-migratory effects of palmitic acid. Using first trimester placental explants, palmitic acid exposure decreased EVT invasion through Matrigel. Our findings reveal that palmitic acid induces an inflammatory response in EVTs and attenuates their migration through a mechanism involving PAI1. High levels of palmitic acid in pathophysiological situations like obesity may impair early placental development and predispose to placental dysfunction.

## Introduction

Extravillous trophoblast (EVT) migration is a critical component of human placentation. EVTs migrate into the decidua as far as the inner third of the myometrium, anchor the placenta to the uterus, integrate into various uterine structures (including endometrial glands and blood vessels) and transform the tissue architecture of the maternal-placental interface [1]. Migrating EVTs displace smooth muscle cells (SMCs) surrounding uterine spiral arteries [2,3], transforming these vessels into low-resistance conduits capable of providing the placenta with a consistent supply of maternal blood that gently bathes the delicate surfaces of the chorionic villi. Defects in EVT migration cause shallow spiral artery remodeling, and are linked to serious pregnancy complications including preeclampsia, which is a major cause of maternal and fetal sickness and mortality [4].

Maternal obesity is a major risk factor for placental dysfunction and various obstetric complications [5–8]. A study evaluating depth of spiral artery remodeling in stillbirths found that elevated body mass index (BMI) was the only maternal characteristic that significantly associated with poor spiral artery remodeling [9]. In rats, a species that, like humans, relies on deep trophoblast invasion for pregnancy success, diet-induced obesity impairs trophoblast migration and spiral artery remodeling, and triggers placental inflammation [10]. Therefore, factors within an obesogenic milieu may impact early aspects of placental development such as EVT migration, and predispose to adverse pregnancy outcomes.

Obesity is associated with elevated plasma levels of free fatty acids [11,12]. In particular, plasma levels of saturated long-chain fatty acids such as palmitic acid are higher in individuals with elevated BMI [13], including pregnant women with elevated pre-pregnancy BMI or excessive gestational weight gain [14]. Palmitic acid is the most common saturated fatty acid in the human body. It has a sixteen-carbon backbone (16:0), and is obtained through dietary intake or synthesized endogenously from other macronutrients. High levels of palmitic acid modulate cellular metabolism and promote production of inflammatory mediators in several cell-types [15–19]. In primary cytotrophoblasts isolated from term placentas, palmitic acid (and stearic acid – another long-chain saturated fatty acid) activates toll-like receptor 4 and stimulates production of pro-inflammatory cytokines including tumor necrosis factor alpha, interleukin (IL)-6, and IL-8 [20]. These effects are not observed in cytotrophoblasts exposed to unsaturated fatty acids, indicating that saturated fatty acids such as palmitic acid may be uniquely capable of promoting inflammation at the maternal-placental interface. In some cell-types, palmitic acid also induces expression of plasminogen activator inhibitor-1 (PAI1) [21], a serine protease inhibitor and powerful regulator of hemostasis, fibrinogenesis and cell migration. PAI1 inhibits EVT motility in vitro, and levels are elevated in pregnancy complications characterized by deficient placentation (e.g. preeclampsia and unexplained recurrent pregnancy loss)[22]. Since excessive inflammation and production of PAI1 is associated with compromised trophoblast function and predisposes to poor placentation [23–26], herein we hypothesize that palmitic acid stimulates expression of PAI1 and other inflammatory mediators in EVTs, and reduces their migratory potential.

In this study, we investigated the effect of SMCs on EVT migration, using co-cultures of vascular SMCs with a well-established EVT-like cell-line as a model system. Next, we determined whether palmitic acid affects migration of EVTs toward SMCs, and we profiled expression of inflammatory mediators, including PAI1, by EVTs following exposure to palmitic acid. We found that palmitic acid stimulates inflammatory pathways in EVTs and impairs their migratory potential. Moreover, we identified PAI1 as a central mediator of the anti-migratory effects of palmitic acid. Our results suggest that high levels of palmitic acid in susceptible individuals may contribute to a suboptimal maternal-placental interface and predispose to deficient placentation.

## Methods

### Cells

HTR8/SVneo cells (henceforth called HTR8 EVTs), a well-established human first trimester EVT-like cell-line derived from placental explant outgrowths [27], were maintained in standard culture conditions (37°C, 5% CO_2_) with RPMI-1640 medium containing 5% fetal bovine serum (FBS), 100 units/ml penicillin, and 100 μM streptomycin (Sigma-Aldrich) for no more than twenty sequential passages. The number of viable HTR8 EVTs was assessed by staining with trypan blue and counting with a hemocytometer.

Primary SMCs derived from human aorta (Cell Applications 354-05a) were maintained in proprietary growth media (Cell Applications 311-500). Cells were plated at a density of 1.5 × 10^4^ cells/cm^2^, and grown in standard culture conditions for up to 15 passages. To differentiate cells to a contractile phenotype, 1.0 × 10^4^ SMCs/cm^2^ were plated, and growth medium was replaced with SMC differentiation medium (Cell Applications 311D-250) for up to seven days. To produce SMC conditioned media, SMCs were differentiated for 5 days, then new differentiation medium was provided and conditioned for 48 h. Conditioned media were then removed, centrifuged, and used for experiments.

Primary human uterine microvascular endothelial cells (PromoCell C-12295) were maintained in proprietary growth media (PromoCell C-22020). Cells were plated at a density of 2.0 × 10^4^ cells/cm^2^ and grown in standard culture conditions for up to 10 passages.

Human embryonic kidney (HEK)-293T cells were maintained in standard culture conditions with Dulbecco’s Modified Eagle’s Medium (DMEM) containing 10% FBS, 100 units/ml penicillin, and 100 µM streptomycin for up to twenty passages.

### Tissue collection

Informed written consent was obtained from each patient in accordance with the Declaration of Helsinki. Collections were approved by the Morgentaler Clinic and the Mount Sinai Hospital Research Ethics Board (Toronto, Canada; REB12-0007E).

For collection of plasma, blood samples were procured from healthy non-pregnant women, pregnant women during mid-second trimester (15.2-17.2 weeks), and third-trimester pregnant women with or without early-onset preeclampsia. Clinical measures of patients are provided in Table 1. Blood samples were collected in sterile tubes containing ethylenediaminetetraacetic acid-dipotassium salt, and plasma was immediately separated from peripheral blood mononuclear cells and polymorphonuclear leukocytes using a dual density gradient separation kit (Histopaque 1119/1077, Sigma-Aldrich), according to the manufacturer’s protocol. Plasma was aliquoted and stored at −80°C until use.

**Table 1.** Clinical measures of patients used for quantification of PAI1 levels in plasma.

| **Demographic** | **Non-Pregnant**<br>**(n=6)** | **Second**<br>**Trimester**<br>**(n=10)** | **Third Trimester**<br>**No Preeclampsia**<br>**(n=9)** | **Third Trimester**<br>**Preeclampsia**<br>**(n=8)** |
| --- | --- | --- | --- | --- |
| **Age (years)**<br>Mean<br>Range | 32.43<br>25-37 | 32.52<br>20 - 42 | 29.86<br>23 - 40 | 26.88<br>25 - 35 |
| **Parity**<br>Nullipara<br>Multipara | 3<br>3 | 5<br>5 | 5<br>3 | 6<br>2 |
| **Gestational age at blood**<br>**collection (weeks)**<br>Mean<br>Range | N/A | 16.33<br>15.2-17.2 | 27.80<br>26.0-29.0 | 29.53<br>27-34.0 |
| **Gestational age at delivery**<br>**(weeks)**<br>Mean<br>Range | N/A | 38.07<br>37 - 41.1 | 39.00<br>37.3 - 40.6 | 31.20*<br>28.2 – 34.4 |
| **Max systolic blood pressure** | N/A | 116.6 ± 11.93 | 124.2 ± 19.83 | 154.79 ± 6.24* |
| **Max diastolic blood pressure** | N/A | 71 ± 7.65 | 72.1 ± 9.89 | 100.33 ± 7.72* |

(\*; P <0.05)

First trimester (5-8 week) placentas, obtained at the time of elective terminations of pregnancy, were used to prepare placental explants as previously described [28]. Villous explants were dissected, washed in PBS, and embedded on Matrigel-coated culture inserts (0.4-μm pores, 12-mm diameter; EMD Millipore). Inserts were placed into wells containing serum-free DMEM/F12 media containing 100 U/ml penicillin/streptomycin, 2 mM L-glutamine, 100 μg/ml gentamicin, and 2.5 μg/ml fungizone, and were incubated at 37°C with an atmosphere containing 3% O_2_ and 5% CO_2_. 48-h following plating, adherent explants that initiated EVT outgrowths were given SMC-conditioned media containing either BSA or 125 μM palmitic acid for 72 h. Outgrowth area was measured using Image J software [29]. Each treatment was conducted in triplicate, and repeated using explants from six different placentas. To determine depth of EVT invasion, the Matrigel plug (containing invaded EVTs) was fixed in 4% paraformaldehyde, embedded in paraffin, and 5-μm serial sections prepared. Sections were rehydrated and stained using hematoxylin and eosin to identify invading EVTs. The depth of EVT invasion into Matrigel was then calculated based on the number of consecutive sections in which EVTs were detected.

### Treatments

To prepare fatty acids, 30% fatty acid-free bovine serum albumin (BSA) was conjugated 2:1 to either 20 mM palmitic acid or oleic acid, and added to SMC conditioned media at final concentrations of 125, 250, and 500 μM (Sigma-Aldrich). Controls consisted of medium containing an equivalent amount of BSA. Activity of P38-MAPK was inhibited using 10 μM SB203580 (P38-MAPK inhibitor); activity of ERK1/2 was inhibited using 10 μM U0126 (MEK inhibitor). Both inhibitors were dissolved in dimethyl sulfoxide (DMSO). SMC-conditioned media containing DMSO was used as control for these experiments.

### Immunofluorescence

Cells were fixed with 4% paraformaldehyde, permeabilized using 0.3% Triton X-100, blocked in 10% normal goat serum (ThermoFisher Scientific), and immersed in antibodies targeting α-smooth muscle actin (A2547, 1:400, Sigma Aldrich), calponin (D8L2T, 1:50, Cell Signaling Technology), or transgelin (sc-53932, 1:50, Santa Cruz Biotechnology). Cells were then incubated with species-appropriate fluorescent antibodies (AlexaFluor, ThermoFisher Scientific), and nuclei counterstained using 4’, 6-diamidino-2-phenylindole (DAPI, ThermoFisher Scientific). Cells were imaged using a Zeiss Axio fluorescence microscope.

### 5-ethynyl-2’-deoxyuridine incorporation assay

5 × 10^4^ HTR8 EVTs were seeded onto Poly-D lysine-coated coverslips. The following day, 10 μM 5-ethynyl-2’-deoxyuridine (EdU, dissolved in culture media) was added to cells for 4 h. Cells were fixed using 4% paraformaldehyde and detection of EdU was performed according to the manufacturer’s instructions (ClickiT EdU Proliferation Kit, ThermoFisher Scientific). Nuclei were detected using Hoechst. Cells were imaged with a Zeiss Axio fluorescence microscope. The total number of cells and number of EdU-positive cells were counted in three random non-overlapping fields of view per well, and percentage of EdU-positive cells was calculated.

### Cell viability and apoptosis assays

HTR8 EVTs were seeded at 5 × 10^4^ cells/cm^2^. The following day, SMC conditioned media containing BSA, palmitic acid, or oleic acid were added for 24 h. As a positive control, 10 μM camptothecin (Cell Signaling Technology) was added to cells for 24 h. Cells were then trypsinized, centrifuged, and incubated with annexin V and propidium iodide (PI) as per the manufacturer’s instructions (Early Apoptosis Detection Kit, Cell Signaling Technology). Cells were analyzed by flow cytometry using a BD FACSCanto cell analyzer (BD Biosciences). Data were analyzed using FlowJo software. Gating strategies can be found in Supplemental Figure 1. In brief, gating commenced with a forward scatter area versus forward scatter height plot to remove doublets, followed by a second gate on forward scatter area versus side scatter area to remove obvious debris from the plot while retaining both live and dead cells. Single-stained positive control cells on annexin V versus PI plots were gated on this population, which was also used to derive the frequencies shown in Figure 3.

**Figure 1.**
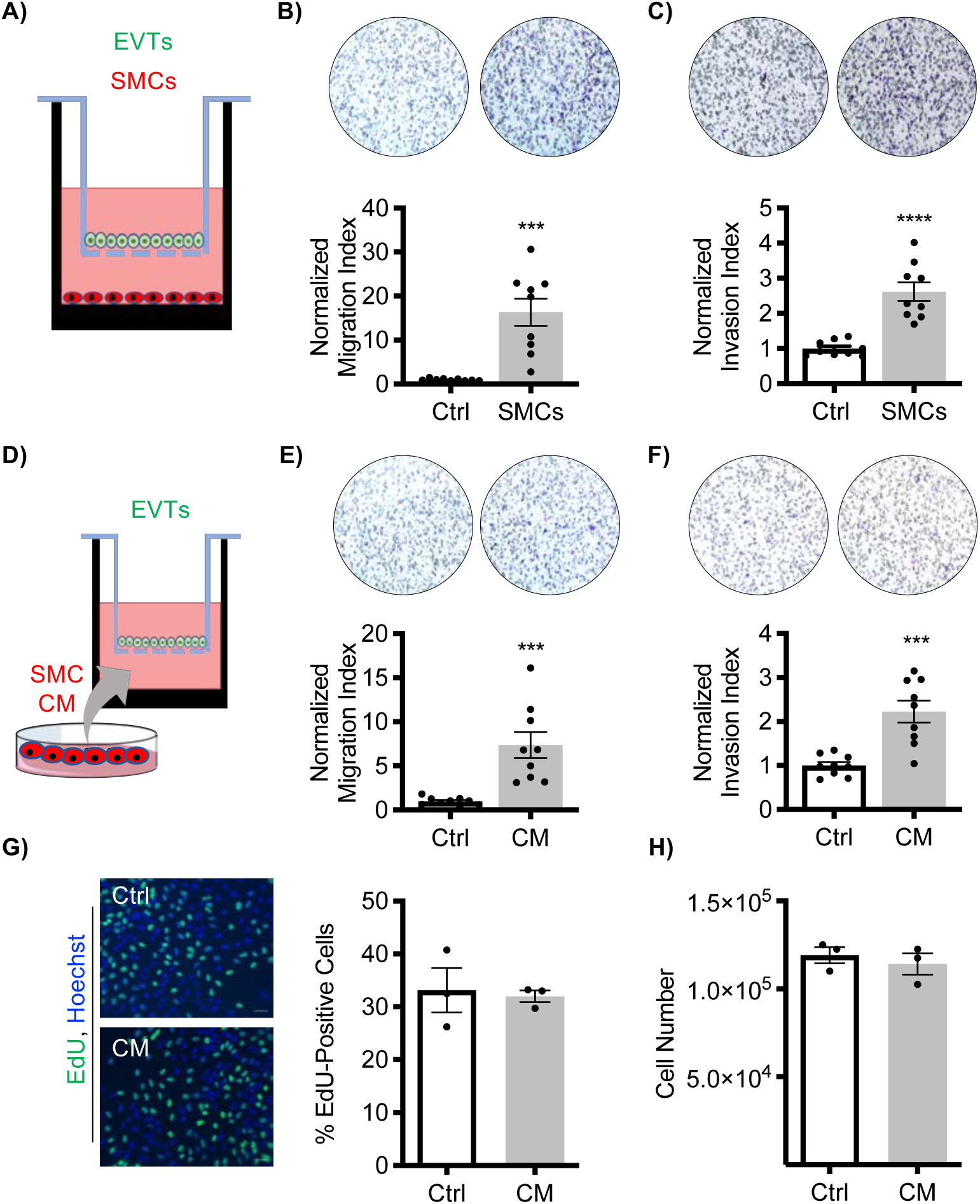
Contractile SMCs enhance migration and invasion of EVTs. **(A)** Schematic of co-culture design with SMCs. Relative number of HTR8 EVTs that migrated **(B)** and invaded **(C)** in the presence of SMCs. Controls (Ctrl) consisted of wells not containing SMCs. Representative images of membranes are included above each graph (the black circles represent pores within the transwell membrane; cells appear purple). **(D)** Schematic of experimental design, showing transwells containing HTR8 EVTs placed into wells containing SMC conditioned media (CM). Relative number of HTR8 EVTs that migrated **(E)** and invaded **(F)** in the presence of SMC CM. Ctrl represents cells migrating toward unconditioned media. Representative images of membranes are included above each graph. **(G)** Percentage of EdU-positive HTR8 EVTs after exposure to SMC CM in comparison to cells immersed in unconditioned media (Ctrl). Representative images are shown to the left of the graph. Nuclei were detected using Hoechst. **(H)** Trypan blue viability assay showing total live cell counts of HTR8 EVTs exposed to SMC CM compared to Ctrl (unconditioned media). Graphs represent means ± SEM. Migration and invasion assays were conducted using 3 membranes per treatment from each of 3 independent experiments. Proliferation and viability experiments: N=3. Asterisks denote statistical significance (***, P<0.001; ****, P<0.0001). Scale bar = 100 µM.

**Figure 2.**
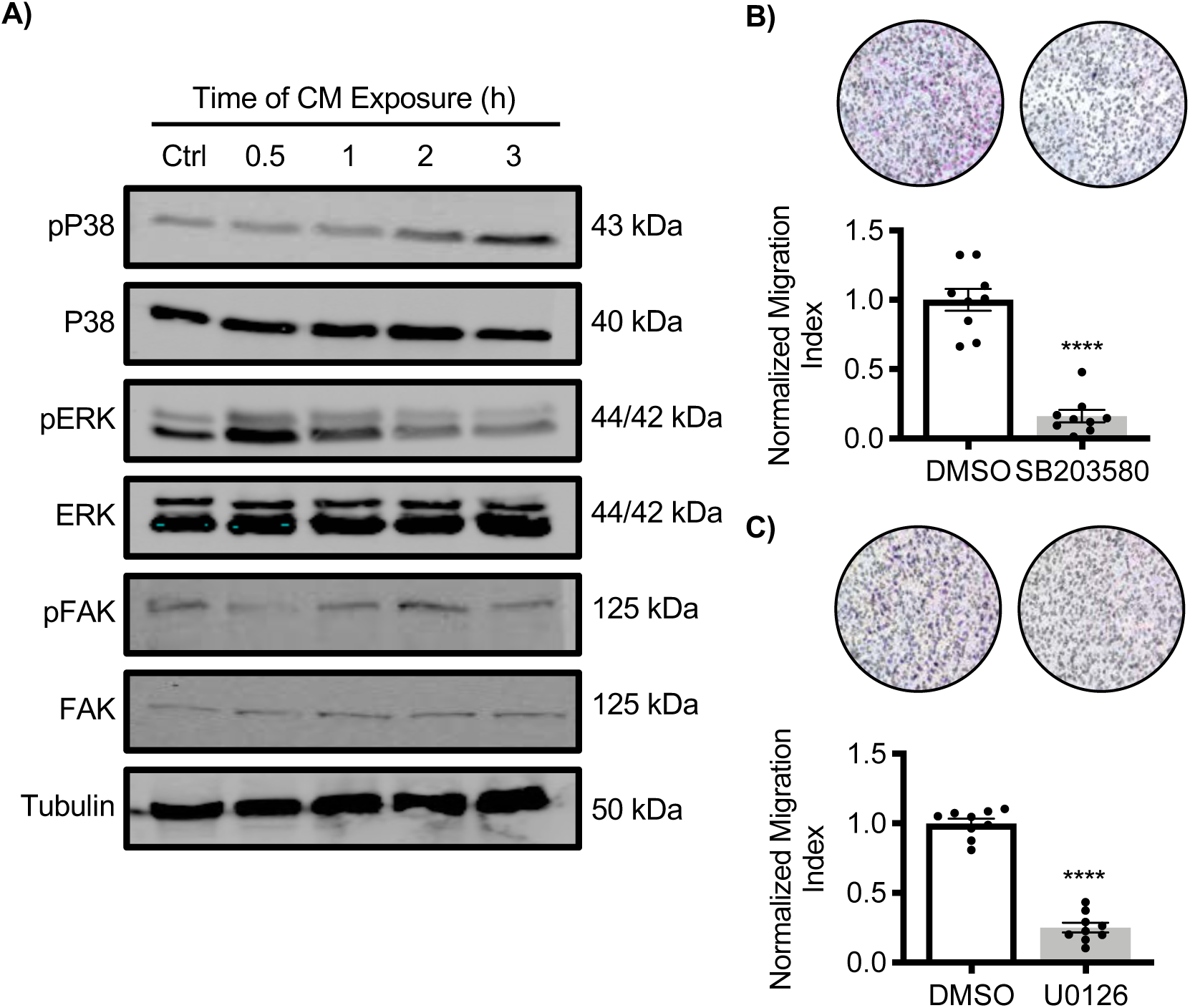
SMCs induce phosphorylation of P38-MAPK and ERK1/2 in EVTs. **(A)** Western blot depicting phosphorylated and total levels of P38-MAPK, ERK1/2, and FAK in HTR8 EVTs following exposure to SMC conditioned medium (CM) for 0.5, 1, 2 or 3 h. Cells not exposed to CM were used as a control (Ctrl). ⍰-tubulin was used as a loading control. Uncropped images of the western blots are provided in Supplemental Figure 5. Relative number of HTR8 EVTs that migrated toward SMC CM containing **(B)** SB203580 (P38-MAPK inhibitor) or **(C)** U0126 (MEK inhibitor) in comparison to media containing DMSO. Representative images of membranes are shown above each graph (the black circles represent pores within the transwell membrane; cells appear purple). Graphs represent means ± SEM. Western blots: N=3 independent experiments; migration assays were conducted using 3 membranes per treatment from each of 3 independent experiments. Asterisks denote statistical significance (****, P<0.0001).

**Figure 3.**
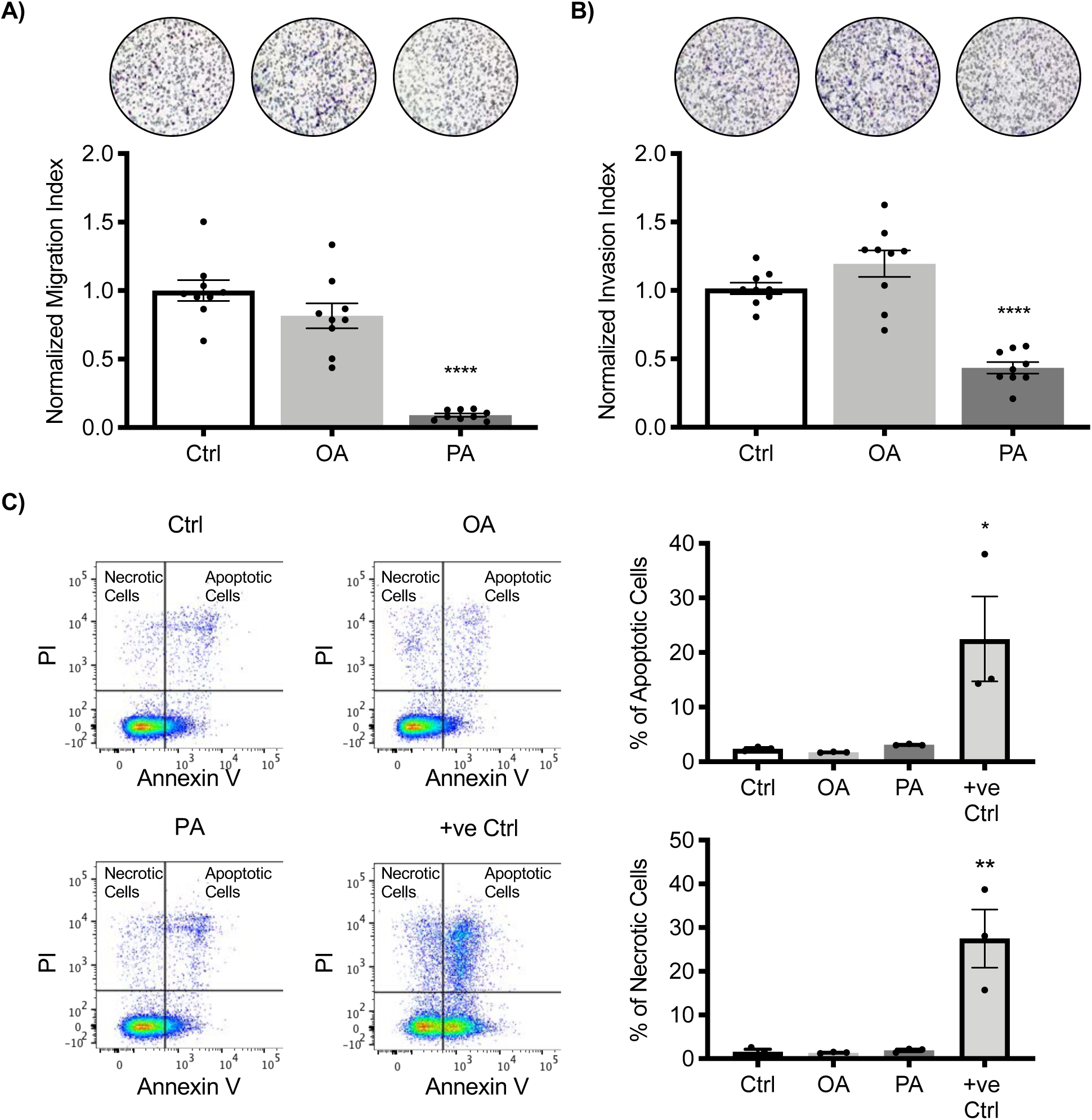
Palmitic acid attenuates SMC-induced EVT migration and invasion. Relative number of HTR8 EVTs that **(A)** migrated or **(B)** invaded through Matrigel in the presence of SMC conditioned media containing either 125 µM oleic acid (OA) or 125 µM palmitic acid (PA). Controls (Ctrl) consisted of cells migrating or invading toward SMC conditioned media containing BSA. Representative images of membranes are included above each graph (the black circles represent pores within the transwell membrane; cells appear purple). **(C)** Flow cytometry analysis of annexin V and PI-positive (apoptotic) and PI-positive (necrotic) HTR8 EVTs treated with BSA (Ctrl), OA, PA, or camptothecin (+ve Ctrl). Percentage of apoptotic and necrotic cells are shown to the right of the images. Gating strategies and single-stained controls can be found in Supplemental Figure 1. Graphs represent means ± SEM. Migration and invasion assays were conducted using 3 membranes per treatment from each of 3 independent experiments. Flow cytometry: N=3. Asterisks denote statistical significance (*, P<0.05; **, P<0.01; ****, P<0.0001).

### Transwell migration and invasion assays

To measure cell migration, 2.0 × 10^4^ HTR8 EVTs were placed into transwell inserts (8-μm pore, 6.5-mm diameter, Greiner BioOne). Media containing 3.8 × 10^4^ human uterine microvascular endothelial cells, 1.9 × 10^4^ synthetic SMCs, 1.9 × 10^4^ contractile SMCs, or conditioned media from contractile SMCs containing either fatty acids or inhibitors (see cell treatments) were added to the lower chamber. Cells were incubated in standard culture conditions for 24 h. Cells in the upper portion of the transwell were removed with a cotton swab. Cells attached to the underside of the membrane were fixed in methanol, then stained with Diff-Quik cytometry stain (GE Healthcare). Membranes were excised, placed onto slides, and counted using a bright-field microscope. Normalized migration indices were calculated by dividing the number of cells that migrated under both control and treatment conditions by the mean number of cells that migrated in control conditions, as done previously [30]. This normalization step was performed to determine the relative change in cell migration for each experiment, which facilitated comparisons between experiments.

Cell invasion was assessed by precoating transwells with growth factor-reduced Matrigel (BD Biosciences, 400 µg/ml diluted in serum-free RPMI-1640) for 3 h. Medium was removed, and then 4.0 × 10^4^ HTR8 EVTs were placed on top of the Matrigel. All other steps were the same as described above. Normalized invasion indices were determined by dividing the number of cells that invaded under both control and treatment conditions by the mean number of cells that invaded in control conditions.

### Immunohistochemistry

Explants were fixed in 4% paraformaldehyde, embedded in paraffin, and sectioned at 5-μm thickness. Sections were deparaffinized, rehydrated, blocked, and immersed in primary antibodies specific for proliferating cell nuclear antigen (PCNA; sc-56, 1:200, Santa Cruz Biotechnology) or human leukocyte antigen-G (HLAG; sc-21799, 1:50, Santa Cruz Biotechnology). The following day, species-appropriate fluorescent antibodies were applied (AlexaFluor, ThermoFisher Scientific), and DAPI was used to counterstain nuclei. Sections were mounted with Fluoromount-G (SouthernBiotech), and images acquired using a Nikon DS-Qi2 microscope.

### Western blot analysis

Cells were lysed using 1× Laemmli sample buffer (2% sodium dodecyl sulfate, 10% glycerol, 5% 2-mercaptoethanol, 0.002% bromophenol, 0.125 M Tris-HCl and 0.5 M dithiothreitol) supplemented with phenylmethylsulfonyl fluoride, boiled, and loaded onto sodium dodecyl sulfate-containing polyacrylamide gels. Proteins were separated using gel electrophoresis, and transferred to polyvinylidene difluoride membranes (GE Healthcare). Membranes were blocked with 3% BSA in TBS containing 0.1% Tween-20, and probed using antibodies targeting α-smooth muscle actin (A2547, 1:2000, Sigma Aldrich), calponin (D8L2T, 1:1000, Cell Signaling Technology), transgelin (SC-53932, 1:500, Santa Cruz Biotechnology), P38-MAPK (D13E1, 1:1000, Cell Signaling Technology), phosphorylated P38-MAPK (D3F9, 1:1000, Cell Signaling Technology), ERK1/2 (9102, 1:1000, Cell Signaling Technology), phosphorylated ERK1/2 (E10, 1:2000, Cell Signaling Technology), focal adhesion kinase (FAK; D507U, 1:1000, Cell Signaling Technology), and phosphorylated FAK (8556, 1:1000, Cell Signaling Technology). Expression of α-tubulin (CP06, 1:1000, Calbiochem) was used as a loading control. Proteins were detected using a LI-COR Odyssey imaging system (LI-COR Biosciences) following incubation with species-appropriate, infrared-conjugated secondary antibodies (Cell Signaling Technology).

### Taqman PCR array and quantitative RT-PCR

RNA was extracted using TriZol (ThermoFisher Scientific). The aqueous phase was diluted 1:1 with 70% ethanol, placed on RNeasy columns (Qiagen), treated with DNase I, and purified. Complementary DNA was generated from purified RNA using High Capacity Reverse Transcription kit (ThermoFisher Scientific), diluted 1:10, and used for quantitative RT-PCR (qRT-PCR). Relative mRNA levels of a panel of inflammation-associated genes were initially screened using a Human Immune Taqman PCR array (4418718, ThermoFisher Scientific) and Taqman Fast Advanced Master Mix. A CFX Connect Real-Time PCR system (Bio-Rad Laboratories) was used for amplification and fluorescence detection. The cycling conditions included an initial uracil-N-glycosylase step (50LJ for 2 min), followed by enzyme inactivation (95°C for 20 s), and then 40 cycles of two-step PCR (95°C for 1 s then 60°C for 20 s). Four reference genes (*18S, GAPDH, HPRT1*, and *GUSB*) were also analyzed in the array, and their expressions were stable among the conditions tested. Transcript levels of select genes (*EDN1, IL6, LIF, IL1A, PTGS2*, and *VEGFA*) were validated on independent samples using qRT-PCR with predesigned Taqman probes (ThermoFisher Scientific, Table 1) and *18S* as a reference RNA. All other qRT-PCR analyses were conducted using cDNA mixed with SensiFAST SYBR Green PCR Master Mix (FroggaBio) and primers described in Table 2. Cycling conditions included an initial holding step (95LJ for 3 min), followed by 40 cycles of two-step PCR (95°C for 10 s, 60°C for 45 s), then a dissociation step (65°C for 5 s, and a sequential increase to 95°C). Relative mRNA expression was calculated using the comparative cycle threshold (ΔΔCt) method, using *18S* as a reference RNA.

**Table 2.**
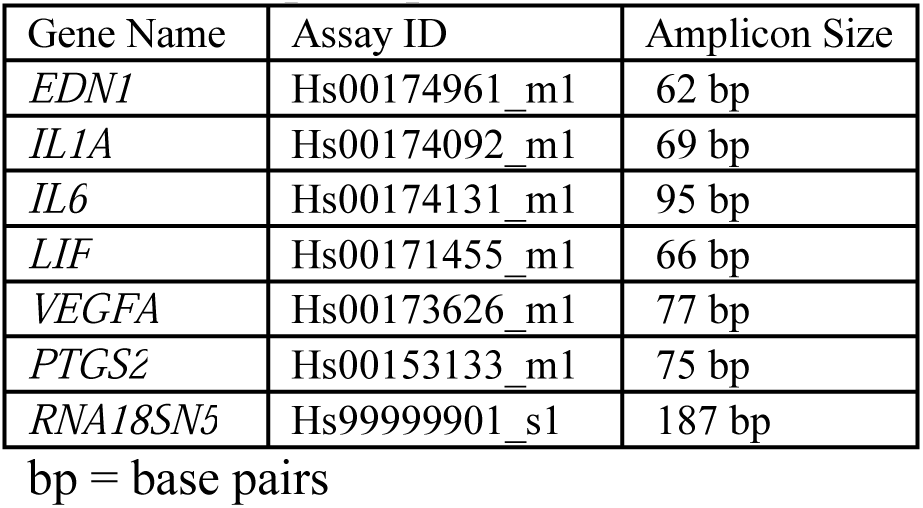
Taqman probe IDs.

| Gene Name | Assay ID | Amplicon Size |
| --- | --- | --- |
| *EDN1* | Hs00174961_m1 | 62 bp |
| *IL1A* | Hs00174092_m1 | 69 bp |
| *IL6* | Hs00174131_m1 | 95 bp |
| *LIF* | Hs00171455_m1 | 66 bp |
| *VEGFA* | Hs00173626_m1 | 77 bp |
| *PTGS2* | Hs00153133_m1 | 75 bp |
| *RNA18SN5* | Hs99999901_s1 | 187 bp |

bp = base pairs

**Table 3.**
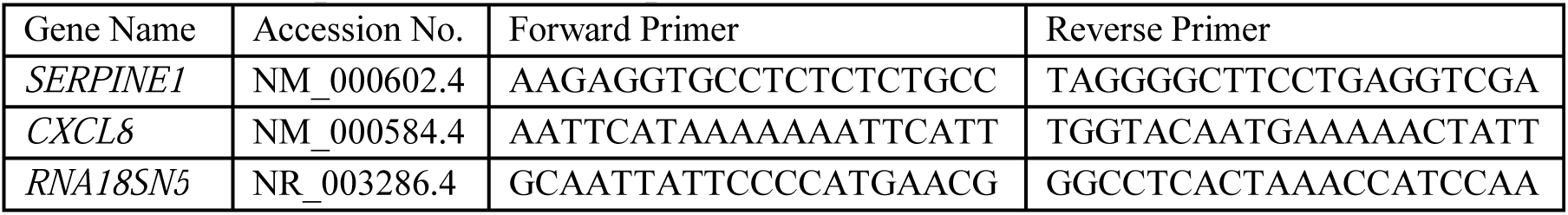
List of primers used for qRT-PCR.

### Enzyme immunoassay

Levels of PAI1 in human plasma were measured using a Bio-Plex multiplex enzyme immunoassay (Customized Human Cancer Biomarker Assay, Bio-Rad Laboratories). Plasma was diluted 1:4 in assay diluent prior to performing the assay. Levels of PAI1 in media conditioned by HTR8 EVTs were measured using a Human Total PAI1 enzyme immunoassay (DY9387-05, Biotechne), as per the manufacturer’s instructions. A standard curve was generated using absorbance values plotted against defined concentrations of recombinant PAI1. Conditioned media were diluted in assay buffer to ensure absorbance values fell within the linear range of the standard curve.

### Lentivirus production

Lentivirus-encapsulated short hairpin RNAs (shRNAs) were generated to knockdown *SERPINE1* gene expression. Briefly, HEK-293T cells were transfected using Lipofectamine 2000 (ThermoFisher Scientific) with lentiviral packaging plasmids (MD2.G, MDLG/RRE, and RSV-Rev) and a *SERPINE1* shRNA construct encoded in a PLKO.1 vector (henceforth called shPAI1: TRCN0000370159, sense: ACACCCTCAGCATGTTCATTG; Sigma-Aldrich). A control PLKO.1 vector containing a scrambled shRNA (Addgene 1864) was used as a control. Lentivirus-containing culture supernatants were collected at 24 h and 48 h, and stored at −80°C until use. To transduce HTR8 EVTs, cells were exposed to lentiviral particles for 24 h in the presence of 8 μg/ml hexadimethrine bromide, as described previously [31]. After 48-h infection, transduced cells were selected with puromycin (3.5 μg/ml).

### Statistical analysis

Statistical comparisons between two means were tested using Student’s t-test; statistical comparisons between three or more means were assessed using analysis of variance, followed by Tukey’s post-hoc analysis. A two-way ANOVA followed by a Sidak’s multiple comparison test was performed to compare HTR8 EVTs transduced with shRNAs and then exposed to BSA or palmitic acid. Means were considered statistically different when P<0.05. GraphPad Prism 7.0 was used for all graphing and statistical analysis. The Human Immune PCR array was conducted using one replicate; all other experiments were repeated at least three independent times. The specific number of replicates is indicated in the figure legends.

## Results

### SMCs stimulate EVT migration and invasion

EVTs migrate chemotactically toward several uterine structures containing vascular SMCs. Therefore, to determine whether SMCs drive migration of EVTs, human vascular SMCs were first differentiated to a contractile phenotype, reminiscent of vascular SMCs surrounding spiral arteries in first trimester human decidua. When cultured under differentiation conditions, vascular SMCs progressively increased levels of calponin, α-smooth muscle actin, and transgelin, and possessed morphological characteristics consistent with a contractile phenotype (Supplemental Figure 2). We then placed uncoated or Matrigel-coated transwells harboring HTR8 EVTs into wells containing differentiated SMCs, to determine whether SMCs drive migration of EVTs (Figure 1A). Compared to control conditions in which SMCs were absent, the presence of SMCs increased HTR8 EVT migration by 16-fold (Figure 1B, P=0.0001) and increased invasion by 2.6-fold (Figure 1C, P<0.0001). Although undifferentiated (synthetic) SMCs and human uterine microvascular endothelial cells enhanced migration of HTR8 EVTs, the extent of migration was not as robust as with differentiated SMCs (3.5-fold and 1.5-fold respectively, P=0.0004 and P=0.0002, Supplemental Figure 3).

To determine whether factors secreted by SMCs stimulated EVT migration, transwells containing HTR8 EVTs were placed into wells containing media conditioned by contractile SMCs (Figure 1D). Compared to unconditioned media, vascular SMC-conditioned media increased HTR8 EVT migration by 7.4-fold (Figure 1E, P=0.0002), and increased invasion through Matrigel by 2.2-fold (Figure 1F, P=0.0005). To investigate whether SMCs affect proliferation of HTR8 EVTs, EdU incorporation and trypan blue viability assays were performed. There was no significant difference in the number of proliferating or viable cells cultured with SMC-conditioned media versus unconditioned media (Figure 1G and H). These results suggest that SMCs drive migration and invasion of HTR8 EVTs, and that this effect is not due to altered cell proliferation.

### SMC conditioned media induces phosphorylation of kinases required for EVT migration

To determine whether contractile SMCs enhance phosphorylation of kinases required for migration of EVTs, we treated HTR8 EVTs with media conditioned by differentiated SMCs, and assessed phosphorylation of signalling factors implicated in EVT migration. There was no change in phosphorylated levels of FAK following exposure of HTR8 EVTs to SMC conditioned media, whereas levels of phosphorylated P38-MAPK and ERK1/2 were increased 3 h and 1 h following exposure to SMC conditioned media, respectively (Figure 2A). Total levels of these kinases were consistent among all treatment conditions. Addition of SB203580 (P38-MAPK inhibitor) to SMC conditioned media resulted in an 84% decrease in HTR8 EVT migration compared to vehicle control (Figure 2B, P<0.0001), whereas treatment with U0126 (MEK inhibitor) inhibited migration by 75% (Figure 2C, P<0.0001). These results suggest that P38-MAPK and ERK1/2 signalling are involved, at least in part, in vascular SMC-induced HTR8 EVT migration.

### Palmitic acid inhibits EVT migration and invasion

Serum levels of palmitic acid are elevated in obese pregnancies, which are associated with poor EVT-directed spiral artery remodeling. Therefore, we next determined whether palmitic acid affects SMC-induced HTR8 EVT migration and invasion. In preliminary experiments, EVT viability was compromised following exposure to 250 μM and 500 μM palmitic acid (not shown), but not following exposure to 125 μM palmitic acid, so a dose of 125 μM palmitic acid was used for all subsequent experiments. Addition of palmitic acid to SMC-conditioned media resulted in a 91% decrease in HTR8 EVT migration and a 57% decrease in HTR8 EVT invasion, compared to cells exposed to BSA (Figure 3, P<0.0001 for both). To determine if these effects were due to the bioactive properties of palmitic acid, we included an additional treatment group in which 125 μM oleic acid (an unsaturated fatty acid) was added to media conditioned by SMCs. Addition of oleic acid had no effect on the capacity of HTR8 EVTs to migrate or invade (Figure 3A and B). To confirm that 125 μM palmitic acid or oleic acid did not affect viability of HTR8 EVTs, annexin V and PI expression were determined by flow cytometry. The number of apoptotic (annexin V and PI-positive) and necrotic (PI-positive) cells did not significantly differ between HTR8 EVTs exposed for 24 h to 125 μM palmitic acid, oleic acid, or BSA. In contrast, exposure of HTR8 EVTs to camptothecin caused a much higher percentage of cells to be apoptotic and necrotic (Figure 3C, P=0.01 and P=0.001). These results demonstrate that palmitic acid impairs SMC-induced migration and invasion of HTR8 EVTs through a mechanism not involving decreased cell viability.

### Palmitic acid induces a pro-inflammatory response in EVTs

Palmitic acid promotes proinflammatory cytokine production in several cell-types [32]. Therefore, to examine whether palmitic acid alters expression of inflammation-associated genes in EVTs, cDNA was prepared from HTR8 EVTs exposed to palmitic acid or BSA, and a PCR array was used to screen inflammation-associated genes potentially induced following palmitic acid exposure. The array included probes for ninety-two distinct genes associated with inflammation, along with four housekeeping genes. Thirty-three inflammation-associated genes were expressed (normalized expression >0.001) in HTR8 EVTs following exposure to either BSA or palmitic acid (Figure 4A). Expression of seven genes (*VEGFA*, encodes vascular endothelial growth factor A; *LIF*, encodes leukemia inhibitory factor; *PTGS2*, encodes cyclooxygenase 2; *END1*, encodes endothelin-1; *IL6*, encodes IL-6; *IL1A*, encodes IL-1α; and *CXCL8*, encodes IL-8) was increased at least 2-fold in EVTs exposed to palmitic acid, so levels of these transcripts were further assessed using qRT-PCR. Palmitic acid increased expression of *VEGFA* (1.6-fold), *EDN1* (2-fold), *IL6* (4.2-fold), and *CXCL8* (11-fold, Figure 4B, all P=0.03, 0.0003, 0.02, and 0.04). Expression of *LIF* also appeared to be increased, although it did not reach statistical significance (4.5-fold, Figure 4B, P=0.056), whereas there was no statistically-significant change in *PTGS2* or *IL1A* (not shown). We additionally investigated expression of *SERPINE1*, because it encodes PAI1: a key component of fibrogenic and thrombotic pathways circulating at increased levels in obese women [33], and in women with early-onset preeclampsia (Figure 5). *SERPINE1* transcript levels in HTR8 EVTs were increased 15-fold following exposure to palmitic acid compared to controls (Figure 4B, P=0.008), and levels of PAI1 in conditioned medium were increased by 62% (Figure 4C, P=0.02). Collectively, palmitic acid enhances expression of various factors associated with inflammation and fibrinogenesis in EVTs.

**Figure 4.**
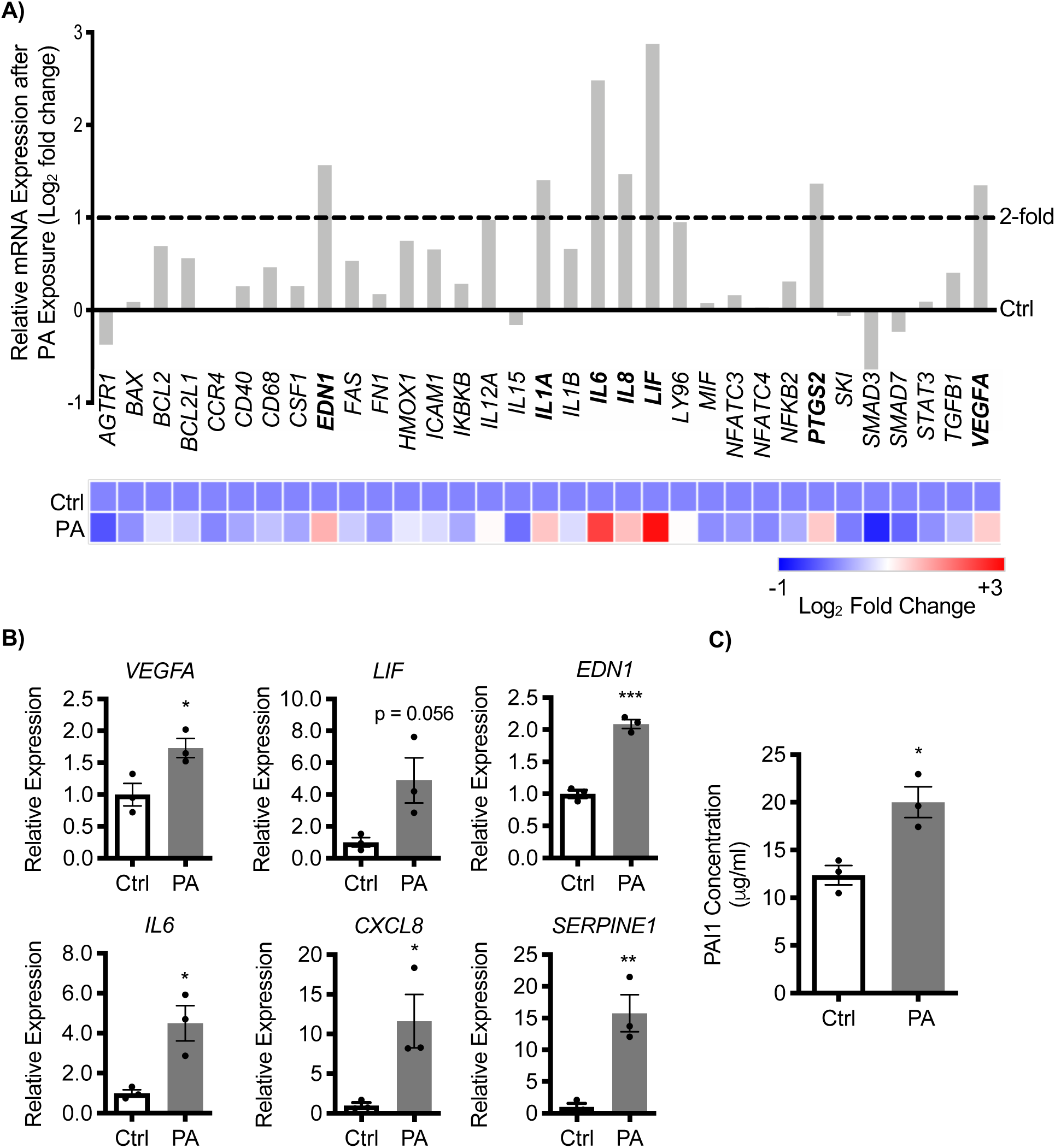
Palmitic acid induces production of inflammatory factors in EVTs. **(A)** PCR array depicting transcript levels of various inflammatory factors following 24-h exposure to SMC conditioned media containing BSA (Ctrl) or 125 µM palmitic acid (PA). Transcript levels from Ctrl conditions are represented by the solid line. Transcripts increased >2-fold (demarcated by the dashed line) following PA exposure are bolded. A heat map is shown below the graph. **(B)** Transcript levels of *VEGFA, LIF, EDN1, IL6, CXCL8*, and *SERPINE1* in HTR8 EVTs after a 24-hour exposure to SMC conditioned media containing BSA (Ctrl) or 125 µM PA. **(C)** Levels of PAI1 in media conditioned by HTR8 EVTs following 24-hour exposure to SMC conditioned media containing BSA (Ctrl) or 125 µM PA. Graphs in **(B)** and **(C)** represent means ± SEM based on N=3 independent experiments. Asterisks denote statistical significance (*, P<0.05; **, P<0.01; ***, P<0.001).

**Figure 5.**
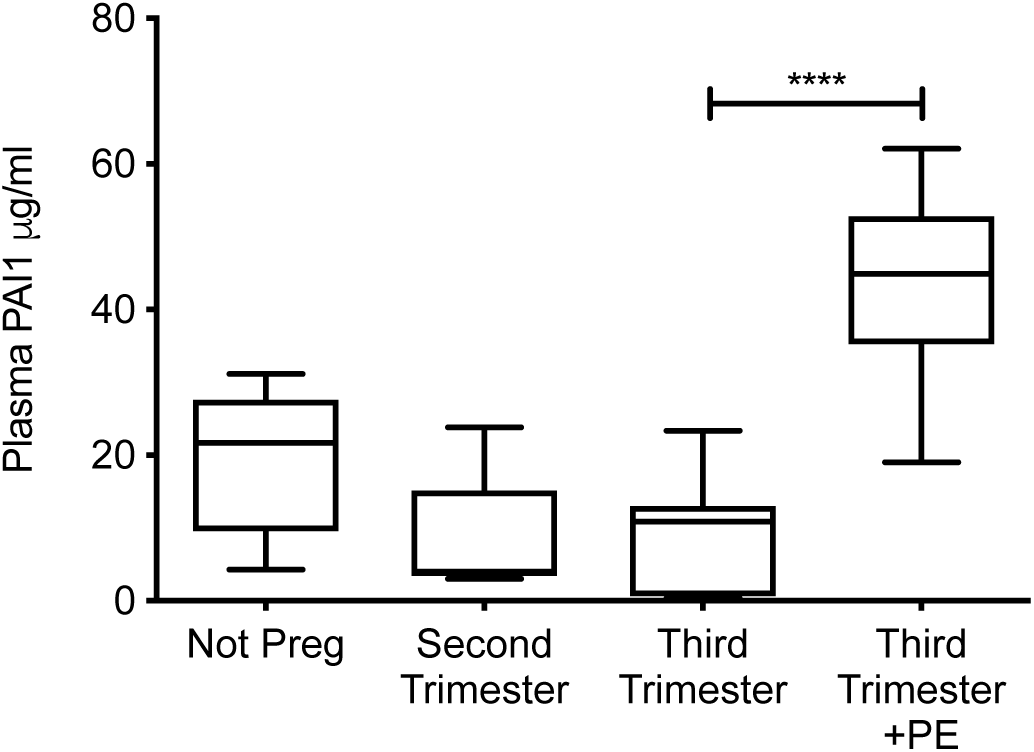
PAI1 is increased in plasma from women with preeclampsia. Plasma was isolated from non-pregnant women (Not Preg; N=6), pregnant women in second trimester (N=10), and pregnant women in early third trimester without preeclampsia (N=9) or with early-onset preeclampsia (PE; N=8). Levels of PAI1 in plasma were detected using a Bio-Plex Assay. Data are shown as box plots. Asterisks denote statistical significance comparing third trimester samples with and without preeclampsia (****, P<0.0001).

### Knockdown of PAI1 rescues trophoblast migration following exposure to palmitic acid

To determine the contribution of elevated PAI1 levels to impaired EVT migration following palmitic acid exposure, we transduced HTR8 EVTs with shRNAs targeting *SERPINE1* (shPAI1). Compared to cells receiving control shRNA, HTR8 EVTs stably expressing shPAI1 exhibited reduced *SERPINE1* expression by 57%, and decreased PAI1 secretion by 75% (Figure 6A and B, P=0.04 and P=0.006). Cells expressing control shRNA exhibited an 86% reduced migration in the presence of palmitic acid compared to BSA-treated conditions (Figure 6C, P=0.0003), which is consistent with our previous observations. Remarkably, migration was completely restored following palmitic acid exposure in shPAI1-expressing EVTs (Figure 6C, P<0.0001 compared to control shRNA-expressing EVTs exposed to palmitic acid). Our results demonstrate that the anti-migratory functions of palmitic acid on EVTs are due, at least in part, to elevated PAI1 expression.

**Figure 6.**
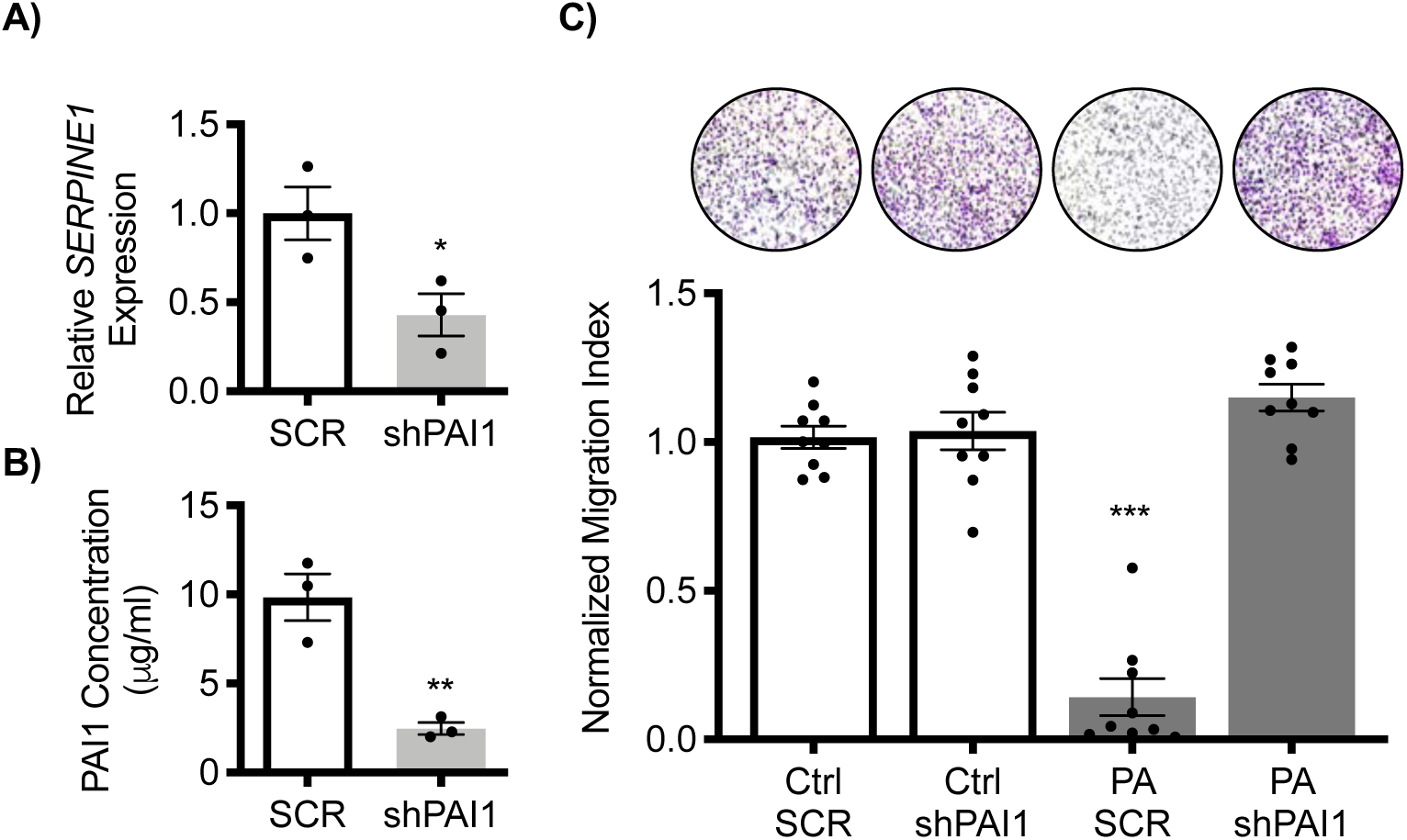
Knockdown of PAI1 restores EVT migration following exposure to palmitic acid. **(A)** Transcript levels of *SERPINE1* in HTR8 EVTs expressing shPAI1 compared to cells expressing control shRNA (scrambled; SCR). **(B)** Levels of PAI1 from media conditioned by HTR8 EVTs expressing SCR or shPAI1. **(C)** Relative number of HTR8 EVTs expressing SCR or shPAI1 that migrated toward SMC conditioned media containing BSA (Ctrl) or palmitic acid (PA). Representative images of membranes are shown above the graph (the black circles represent pores within the transwell membrane; cells appear purple). Graphs represent means ± SEM. **(A)** and **(B)** represent data from 3 independent experiments; **(C)** represents data obtained using 3 membranes per treatment from each of 3 independent experiments. Asterisks denote statistical significance (*, P<0.05; **, P<0.01; ***, P<0.001).

### Palmitic acid inhibits EVT differentiation in placental explants

Primary placental explants were used to further investigate the effects of palmitic acid on EVT cell invasion *ex vivo*. Explants can be used to recapitulate the multiple stages of EVT lineage development, including proliferative PCNA-positive proximal column EVTs and invasive HLAG-positive distal column EVTs (Figure 7A). After 72 h in culture with SMC conditioned media, EVT outgrowth was apparent in all explants, although those exposed to palmitic acid exhibited a 2-fold increased EVT outgrowth area as compared to those exposed to BSA (Figure 7B, P=0.04). However, explants exposed to palmitic acid exhibited a 35% decrease in depth of EVT invasion into Matrigel in comparison to those exposed to BSA (Figure 7C, P=0.01), suggesting that palmitic acid impaired the invasive capacity of distal column EVTs.

**Figure 7.**
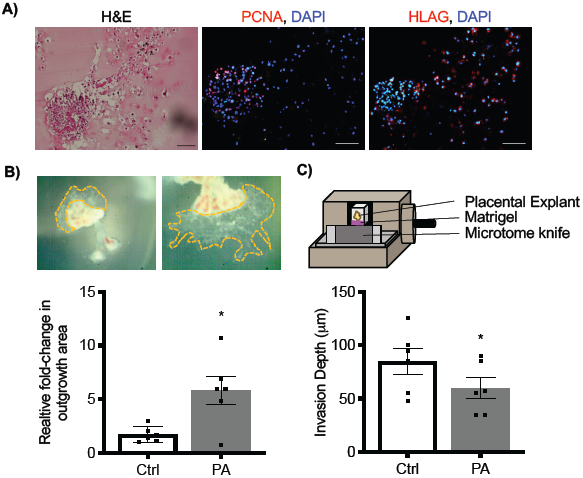
Increased outgrowth and impaired EVT invasion in first trimester placental explants exposed to palmitic acid. **(A)** 8-week placental explants cultured for 96 h stained with hematoxylin and eosin (H&E), PCNA, and HLAG. Nuclei were detected using DAPI. **(B)** Relative change in placental explant outgrowth area after exposure to SMC conditioned media containing BSA (Ctrl) or 125 µM palmitic acid (PA). Representative images of the explants are depicted above the graph. **(C)** Depth of EVT invasion into Matrigel following exposure to SMC conditioned media containing BSA (Ctrl) or PA. Graphs represent means ± SEM based on 3 explants prepared from each of 6 different placentas. Asterisks denote statistical significance (*, P<0.05). Scale bar = 100 µm.

## Discussion

EVT invasion is a critical component of normal placentation and maintenance of a healthy pregnancy. Since EVTs typically migrate toward uterine structures containing smooth muscle (myometrium, uterine glands, endometrial veins, spiral arteries) [1], we first tested the hypothesis that EVTs are driven to migrate toward SMCs. We found that factors produced by contractile vascular SMCs stimulate phosphorylation of ERK1/2 and P38-MAPK in HTR8 EVTs, resulting in enhanced migration and invasion of these cells. We further show that palmitic acid restrains EVT motility and increases expression of several inflammatory factors in EVTs, most notably PAI1, and that the effects of palmitic acid on EVT motility are restored by reducing expression of PAI1. Our results provide new insights into mechanisms of EVT migration, with important implications for pathophysiological conditions such as obesity, which are characterized by elevated levels of palmitic acid and a higher incidence of placental dysfunction.

Although EVTs migrate toward many different uterine structures, the best understood paradigm is their migration toward spiral arteries. EVTs home toward spiral arteries using both interstitial and endovascular routes and transform these vessels by displacing endothelial cells and SMCs. Previous studies have shown that EVTs and EVT cell-lines trigger SMC migration and apoptosis, indicating that EVTs and SMCs are capable of dynamic cellular crosstalk [34– 36]. To the best of our knowledge, our study is the first to show that EVT migration is triggered by factors secreted by contractile vascular SMCs. Although we did not deduce which factors produced by SMCs are responsible for stimulating EVT migration, we did profile a select number of growth factors produced by contractile vascular SMCs (not shown), and detected high expression of platelet-derived growth factor (PDGF), epidermal growth factor (EGF), and heparin-binding EGF-like growth factor (HB-EGF). PDGF, EGF, and HB-EGF stimulate human EVT adhesion, migration and invasion [37–40], so it is possible that production of these growth factors by vascular SMCs contributes to the enhanced EVT migration observed in our study. Regardless of which factors are involved, we show that SMC conditioned media activates ERK1/2 and P38-MAPK signaling pathways in EVTs, and inhibition of either pathway inhibits EVT migration, which is consistent with previous reports [41].

Maternal obesity is a pregestational factor associated with a higher risk of placental dysfunction [42,43]. Pregnant obese women, and those with excessive gestational weight gain, have increased saturated fatty acids circulating in blood, most prominently palmitic acid [14,44]. Palmitic acid concentrations are normally maintained under stringent homeostatic control, likely due to its essential role in cell membrane structural properties, synthesis of palmitoylethanolamide, and protein palmitoylation. Palmitic acid is metabolized in cells into saturated phospholipids (e.g. lysophosphatidylcholine), diacylglycerol, and ceramides, high levels of which can alter cellular signaling events, including oxidative and endoplasmic reticulum stress and activation of protein kinase C. Palmitic acid is also a toll-like receptor agonist, and can stimulate inflammatory responses through myeloid differentiation factor 88/nuclear factor kappa B and interferon regulatory factor 3-dependent pathways (reviewed in [32]). Thus, palmitic acid is a highly bioactive molecule with the potential to alter cellular gene expression, metabolism, and behavior. We found that palmitic acid decreased viability of HTR8 EVTs cells at concentrations of 250 and 500 μM, but cell viability was unaffected at 125 μM, which is why this dose was used for our experiments. Although the concentration of palmitic acid present at the maternal-placental interface during early pregnancy is not known, 125 μM palmitic acid is consistent with other cell culture studies modeling hyperlipidemia seen in obesity and related comorbidities [45–50]. Others have reported compromised trophoblast viability following exposure to high doses of palmitic acid, including endoplasmic reticulum stress, proliferation defects, and reduced viability in HTR8 EVTs exposed to 400 and 800 μM palmitic acid, as well as lipotoxicity in human syncytiotrophoblast exposed to 200 and 400 μM palmitic acid [51,52]. Our study is the first to show the effect of palmitic acid on EVT motility at sublethal doses. Palmitic acid can either stimulate [53–55] or inhibit [56–58] migration of distinct cell-types, suggesting that the impact of palmitic acid on cell motility is likely dose, cell-type, and context-specific. We found that palmitic acid attenuated migration and invasion of HTR8 EVTs, and reduced EVT invasion in first trimester placental explants, whereas there was no effect on EVT motility when cells were exposed to an unsaturated fatty acid, oleic acid. Furthermore, palmitic acid induced expression of several genes encoding inflammatory (*IL6, CXCL8*), vasoactive (*EDN1*), and fibrogenic (*SERPINE1)* proteins, all of which are increased in serum of obese patients [59,60] and in various pregnancy complications [22,61]. High levels of palmitic acid in susceptible pregnancies may therefore contribute to a suboptimal microenvironment at the maternal-placental interface that impairs EVT motility and predisposes to deficient placentation.

In the current study, we detected elevated levels of PAI1 in women with early-onset preeclampsia compared to age-matched controls, which is consistent with several other studies [62–64]. Unfortunately, we did not have patient consent to obtain additional information such as body mass index or palmitic acid concentrations in blood, so correlating these parameters with PAI1 concentrations is the subject of future investigations. High levels of PAI1 may impede placental development by promoting occlusive lesions and deposition of fibrin within placental vasculature [65], as well as restricting EVT migration by inhibiting degradation of extracellular matrices [66–68]. Conversely, low expression of PAI1 is associated with uncontrolled trophoblast invasion in molar pregnancies [66]. In the current study, treatment of HTR8 EVTs with palmitic acid increased PAI1 expression. Palmitic acid induces PAI1 in other cell-types (e.g. renal epithelial cells), indicating that high circulating levels of palmitic acid may enhance PAI1 production by many tissues, resulting in elevated systemic levels of this protein [21]. We found that knocking down expression of PAI1 did not affect migration of EVTs in control conditions, which we attribute to EVTs already migrating at high capacity. Intriguingly, when EVTs were exposed to palmitic acid, motility was abrogated in control cells but completely restored in PAI1-deficient cells. Antibody-mediated neutralization of PAI1 also rescues EVT migration following exposure to the pro-inflammatory cytokine tumor necrosis factor alpha [69]. Increased PAI1 levels may therefore contribute to poor EVT migration and deficient placentation in various pathophysiological conditions, such as in pregnancies with high plasma levels of palmitic acid or inflammatory cytokines.

In sum, our findings show that EVT migration is stimulated by SMCs, and that palmitic acid interferes with this process. We recognize that palmitic acid is only one factor of many circulating at aberrant levels in blood of patients with metabolic disturbances such as obesity that may influence EVT gene expression signatures and migratory phenotypes. However, palmitic acid is the most common saturated fatty acid circulating in human blood [70], and our data show that it is sufficient to induce inflammatory and fibrogenic mediators in EVTs. Thus, altered levels of this particular fatty acid may have major impacts on trophoblast function and placental development. We exploited HTR8 EVTs for analysis of PAI1 expression and knockdown since these cells are advantageous as models of migratory first trimester EVTs and are amenable to stable incorporation of shRNAs. Although technical limitations precluded us from using shRNA to disrupt PAI1 expression in placental explants, in future studies it would be interesting to determine whether neutralization of PAI1 via recently-developed pharmacological inhibitors [71], is sufficient to restore EVT invasion following exposure to palmitic acid. Our findings are in support of others [72], who suggest that monitoring PAI1 levels may have diagnostic utility as a biomarker to predict placental insufficiency. Furthermore, our study opens doors to new therapeutic interventions in which managing levels of palmitic acid or PAI1 to within a “normal” physiological range may help to restore trophoblast function and prevent dysfunctional placentation. Considering the current prevalence of obesity in women of child-bearing age, this intervention may be useful in reducing the high-risk of adverse pregnancy outcomes common to obese pregnancies.

## Acknowledgments

The authors wish to thank Peeyush Lala (University of Western Ontario) for providing HTR8 EVTs; and Kristin Chadwick and Dendra Hillier (University of Western Ontario) for technical support. The authors thank the donors, the Research Centre for Women’s and Infants’ Health BioBank Program at the Lunenfeld-Tanenbaum Research Institute, and the Mount Sinai Hospital and University Hospital Network Department of Obstetrics & Gynecology for the human specimens used in this study.

