## Supplemental Figure Legends for "Palmitic acid induces inflammation in placental trophoblasts and impairs their migration toward smooth muscle cells through plasminogen activator inhibitor-1"

**Supplemental Figure 1. Gating Strategy for Flow Cytometry.** HTR8 EVTs were treated with 10 μM camptothecin for 24 hours. **(A)** Gating used to exclude doublets. **(B)** Gating used to exclude debris. **(C)** PI single-stained control used to acquire gate to identify PI-positive cells. **(D)** Annexin V single-stained control used to acquire gate to identify Annexin V positive cells. **(E)** Annexin V and PI-stained positive control sample.

**Supplemental Figure 2. Differentiation of SMCs to a contractile phenotype.** SMCs were cultured in normal growth media (0 days of differentiation), or SMC differentiation media for up to 7 days. **(A)** Western blot depicting expression of contractile proteins (calponin, α-smooth muscle actin (⍺-SMA), and transgelin) in SMCs cultured for up to 7 days in SMC differentiation media. ⍺-tubulin was used as a loading control. Uncropped images of western blots are provided in Supplemental Figure 4. **(B)** Immunofluorescent images of calponin, ⍺-SMA, and transgelin in SMCs cultured in normal growth media (Control) or 5 days culture in SMC differentiation media. Scale bar = 100 µm.

**Supplemental Figure 3. HTR8 EVT migration is stimulated by undifferentiated (synthetic) SMCs and human uterine microvascular endothelial cells. (A)** Schematic of experimental design. Relative number of HTR8 EVTs that migrated in the presence of (**B**) synthetic SMCs and **(C)** uterine microvascular endothelial cells (ECs). Controls (Ctrl) consisted of wells not containing synthetic SMCs or ECs. Graphs represent means ± SEM. Migration and invasion assays were conducted using 3 membranes per treatment from each of 3 independent experiments. N=3. Asterisks denote statistical significance (***, P<0.001).

**Supplemental Figure 4. Uncropped western blots from Supplemental Figure 2.**

**Supplemental Figure 5. Uncropped western blots from Figure 2A.**
