## Supplemental Figures for "Palmitic acid induces inflammation in placental trophoblasts and impairs their migration toward smooth muscle cells through plasminogen activator inhibitor-1"

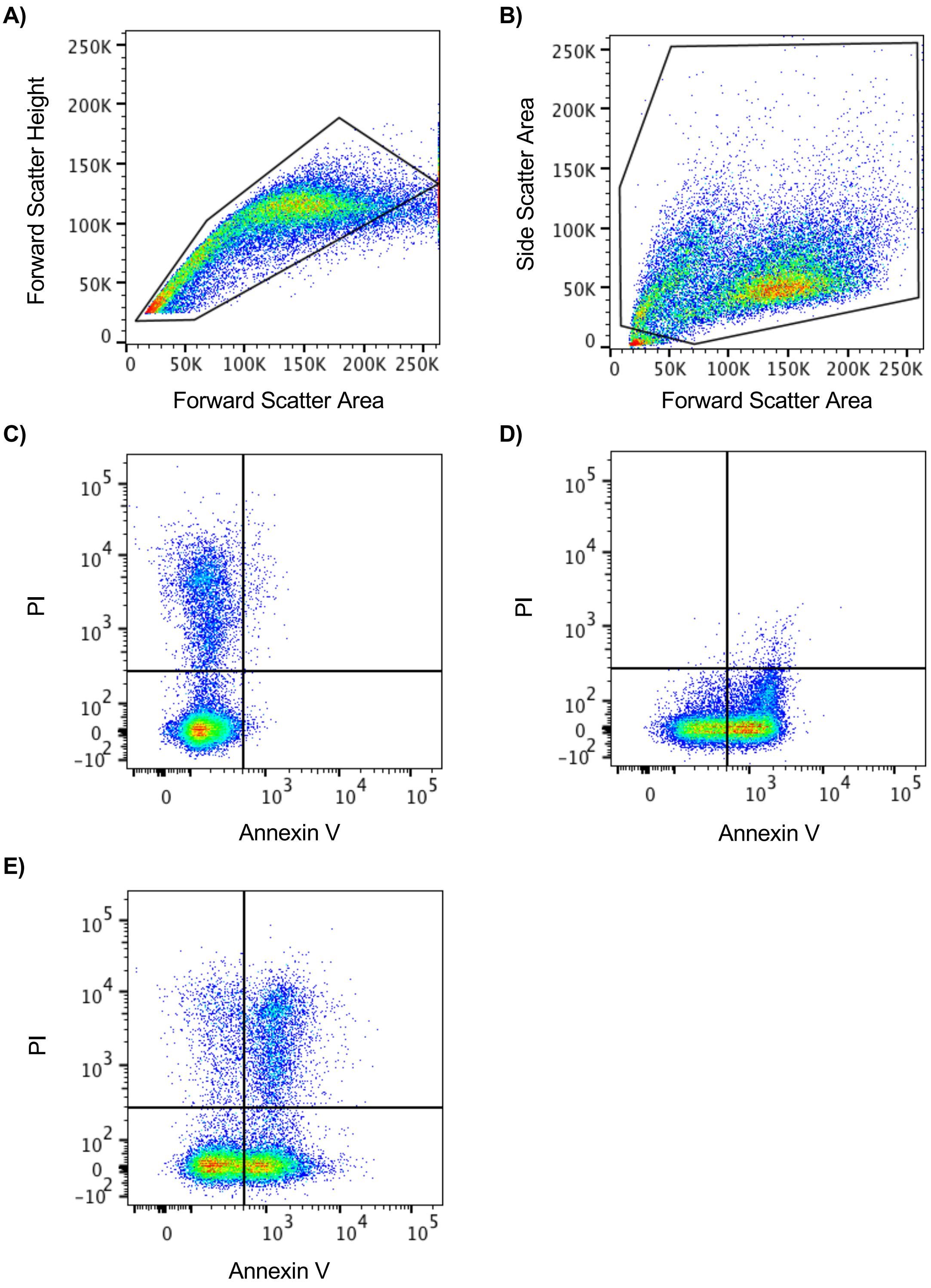


Supplemental Figure 1

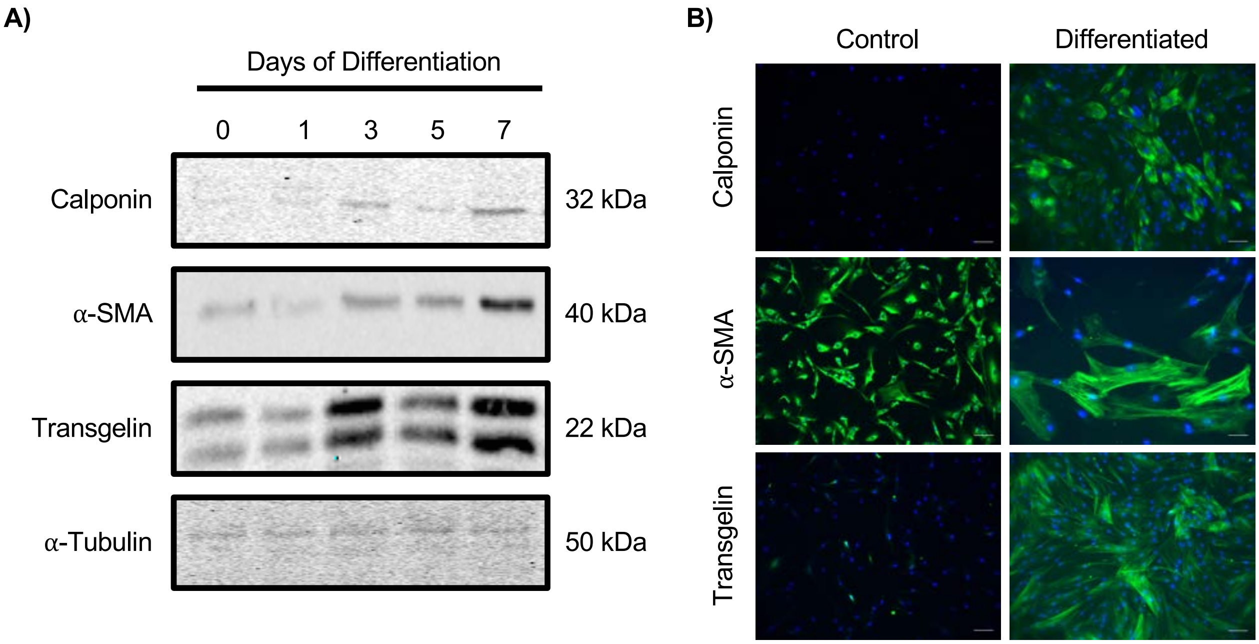


Supplemental Figure 2

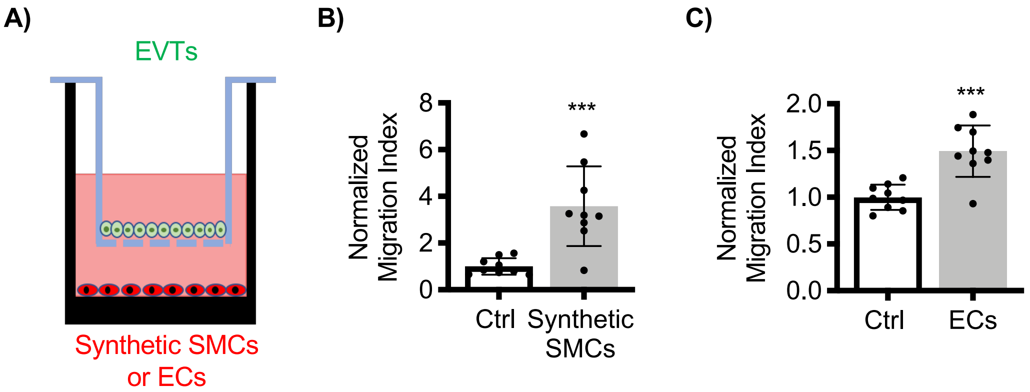


Supplemental Figure 3

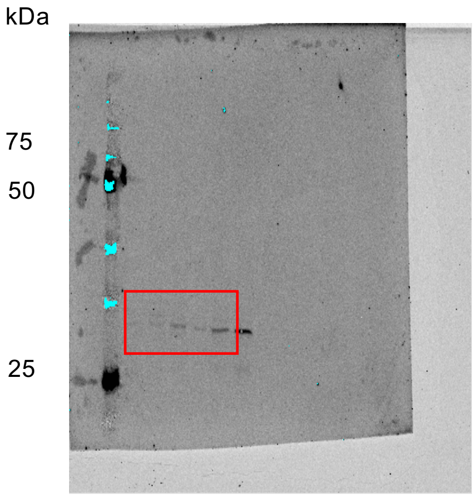


Unprocessed western blot image for Supplemental Figure 1 using antibody: Calponin.

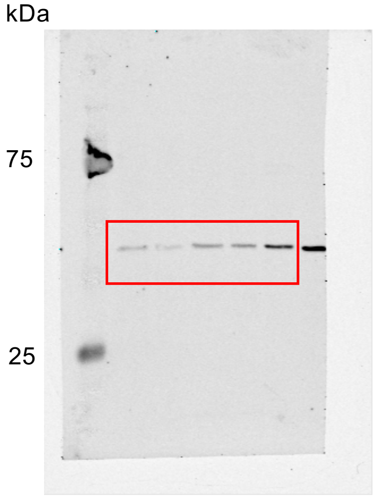


Unprocessed western blot image for Supplemental Figure 1 using antibody: α-smooth muscle actin.

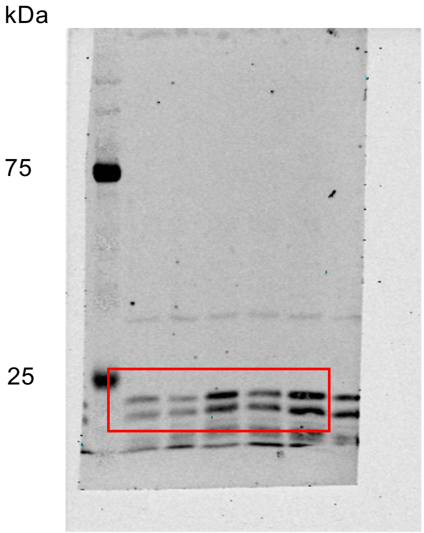


Unprocessed western blot image for Supplemental Figure 1 using antibody: Transgelin. A non-specific band appears at approximately 35 kDa.

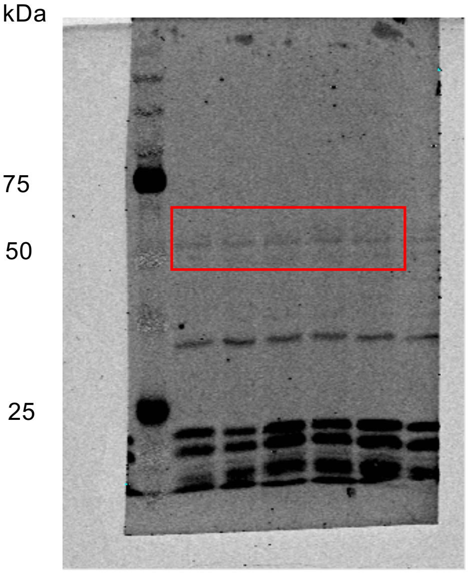


Unprocessed western blot image for Supplemental Figure 1 using antibody: Tubulin. Antibody for Transgelin was also applied to membrane showing bands at approximately 22 kDa. A non-specific band appears at approximately 35 kDa.

Supplemental Figure 4

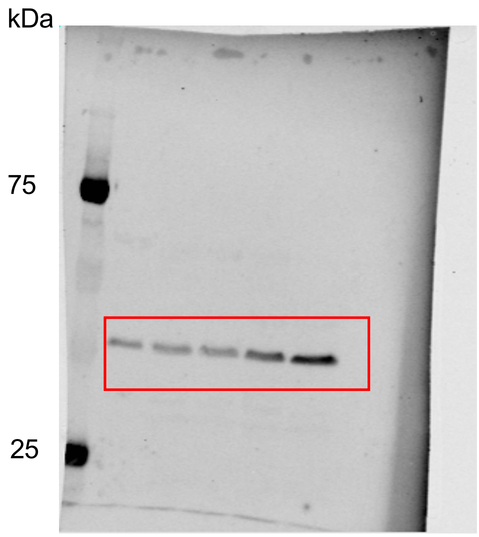

Unprocessed western blot image for Figure 2A using antibody: pP38.

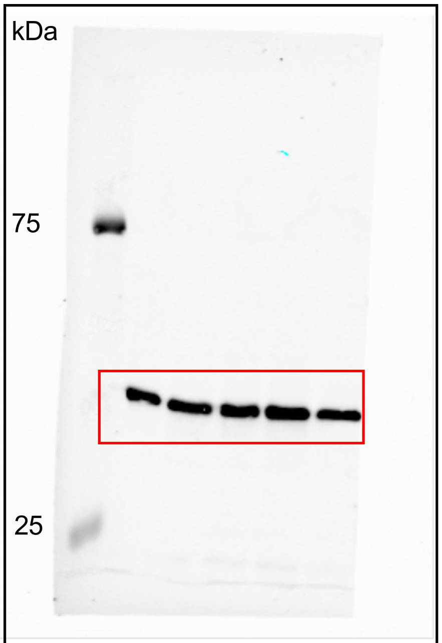

Unprocessed western blot image for Figure 2A using antibody: Total P38.

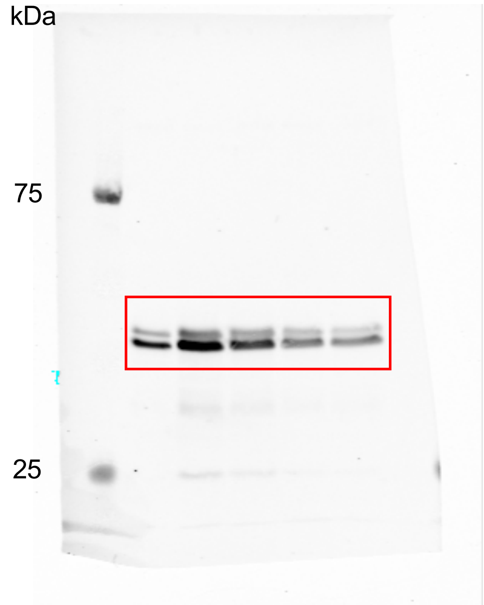

Unprocessed western blot image for Figure 2A using antibody: pERK1/2.

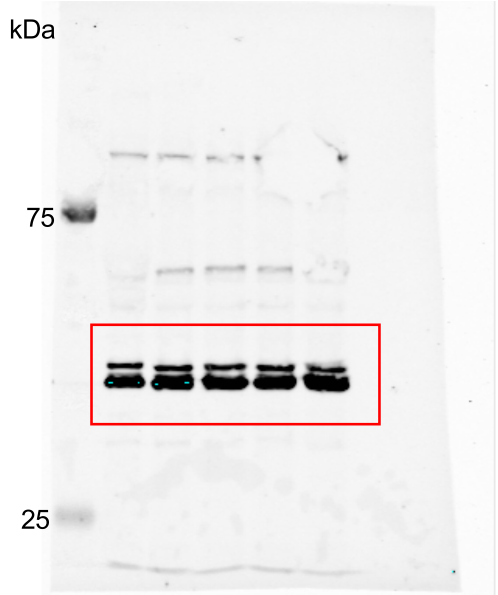

Unprocessed western blot image for Figure 2A using antibody: Total ERK1/2.

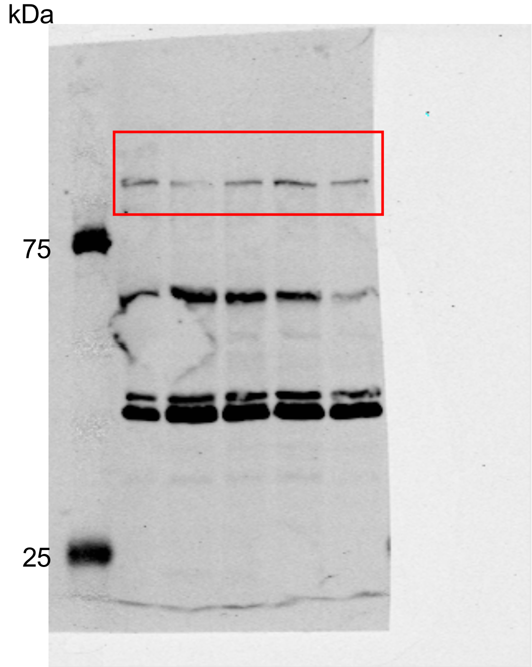

Unprocessed western blot image for Figure 2A using antibody: pFAK. Antibodies for pAKT and total ERK1/2 were also applied to membrane showing bands at 60 kDa and 44/42 kDa, respectively.

kDa

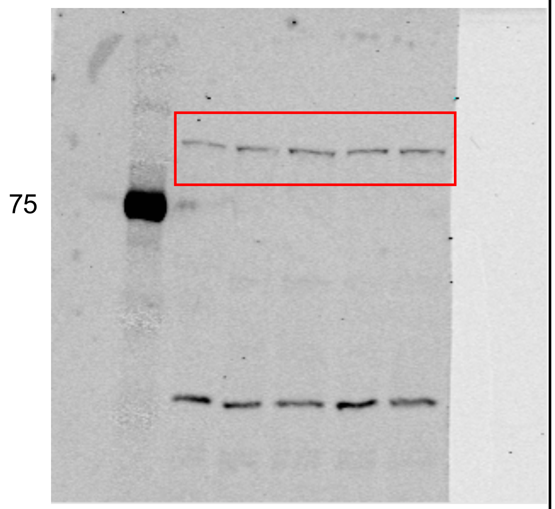


Unprocessed western blot image for Figure 2A using antibody: Total FAK. A non-specific band appears at approximately 35 kDa.

kDa

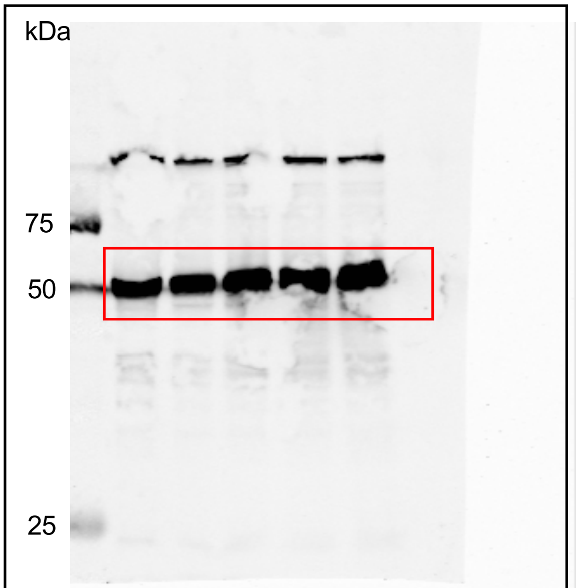


Unprocessed western blot image for Figure 2A using antibody: Tubulin. Antibody for pFAK was also applied to membrane showing a band at 125 kDa.
